# Prognostic Gene Signature for lung adenocarcinoma with a Higher Risk for Treatment Failure and Accelerated cell cycle and cell proliferation Pathway Activity

**DOI:** 10.1101/2023.05.19.540490

**Authors:** Zhen-Qing Li, Zheng Li, Yu-Qi Meng, Xuan Li, Ke-Rong Zhai, Shi-Lin Wei, Bin Li

**Affiliations:** Department of Thoracic Surgery, Lanzhou University Second Hospital, Lanzhou University Second Clinical Medical College, Lanzhou 730030, China; Medical College of Lanzhou University, Lanzhou, China

**Author notes:** Correspondence: Bin Li, Department of Thoracic Surgery, Lanzhou University Second Hospital, Lanzhou University Second Clinical Medical College, Cuiyingmen, Chengguan District, Lanzhou 730030, Gansu Province, China.

## Abstract

Lung cancer is the most common of malignant cancers which represents poor prognosis and high invasive. However, the reliable prognosis genes signature and the differential between the genetic and epigenetic are remain unclear. Here, we confirmed 78 genes of signature A to construct a model to predicted the prognosis and the activity of tumorigenesis-related pathways. The score based on the 78 genes expression is high correlation with the cell cycle and cell proliferation activity. Based on the median of score, the lung adenocarcinoma patients were divided into two cohorts which presents different prognosis, tumorigenesis-related pathways activity. The high score group was enriched in the E2F target, mtorc1 signaling and MYC targets. The low score group can be identified as better prognosis subtype. Also, we used the GDSC data to predicted the PIM inhibitor (AZD1208) as the effective compounds for high score group patients, which is supported by a higher PIM1 and PIM2 expression in the score group. An integrative analysis of multi-omics data identified ECT2 as key nodes in a regulatory network related to the prognostic phenotype. In conclusion, our data identified a molecular classifier for the different score groups patients, and the high score group patients might benefit from treatment with PIM inhibitor.

## Introduction

Because of the high mortality and incidence rate, the lung cancer is the main cause of cancer-related deaths worldwide. And however, the drive mechanism and pathogenic of lung cancer are still unknown. And the main pathogenic as we know is the long-term high-frequency smoking(1). Also, the person who is exposure to air pollution, genetic factors, carcinogen and metabolic activities in long-term time are easily to be lung cancer patients (2). There are two histopathological subtypes, including the small cell lung cancer (SCLC) and non-small cell lung cancer (NSCLC) and the NSCLC constitutes lung adenocarcinoma (LUAD), large cell carcinoma and lung squamous cell carcinoma, which are accounting for 85% of all lung cancer cases (3). Now, the traditional clinical traits such as TMN stage and grade which can help to chose the clinical treatment and predict survival of cancers patients. And different stage and other pathologic factors have different prognosis and the response of treatments were also heterogeneous. Thus, it is urge to find a new target for prognostic prediction. And based on the molecular subtypes, the establish individualized treatment plans can extend the survival of patients. Also, the establish of molecular subtypes of different prognosis may help to deep the understanding of genomic metastasis and development of cancers.

Molecular subtypes have been used to explore NSCLC heterogeneity. Gene expression subtypes of LUSC and LUAD were respectively proposed by the Cancer Genome Atlas (TCGA) research network [5,6]. A recent study also identified a multi-platform-based molecular subtype of NSCLC, including 9 subtypes in 1023 NSCLC patients [7]. There are several other types of lung cancer molecular subtypes based on different gene sets [8,9]. However, early NSCLC has some special molecular characteristics. In these subtypes and gene concentrations, the prognostic information of patients is also not well used, resulting in a weak prognostic ability of patients. However, differences were observed among the molecular subtypes and their prognostic values. This may be due to the limited number of study populations and regional differences.

Here, we used the multi-clustering methods to confirmed the two clusters with different prognosis outcomes. The cluster A was identified as better prognosis outcome and with low activation of cell proliferation pathways. The cluster B with worse prognosis was ascribed to high activation of cell proliferation pathways. And the 78 genes was identified as the cluster B signature gene. Based on those genes, we used the ssgsea algorithm to construct a model which can predict the activity of cell proliferation related pathways and prognosis.

Our study aimed to built a model of prognosis prediction for enabling the stratification of LUAD patients at different activity of cell proliferation and to predict promising drug targets for a more effective and less toxic therapy. Meanwhile, we conducted an integrative analysis of multi-omics data to find key gene of a functional network and based on the GDSC data, our study proposed potential drug targets for high activity of cell proliferation.

## Materials and Methods

### Datasets and Samples

We download 4 GEO datas including GSE31210, GSE30219, GSE68465 and GSE72094 which had detailed survival information and TCGA-LUAD was obtained from TCGA database with detail clinical information. The “Combat” packages was used to reduce probable batch effects. Meanwhile, we used the GSE50081 and GSE37745 to verify the accuracy of model.

### Gene set variation analysis (GSVA)

We used the 50 hallmark pathways from molecular signature database to conduct pathway analyses which was exported from the GSEA base package. For reducing the overlaps and redundancies of pathways, we trimmed the gene sets associated with pathways to keep unique genes of each pathways. And then the GSVA package was used to estimate the pathways score, as implemented in the GSVA package(27) (version 1.22.4).

### Multi omics datas clustering and subtype identification

The two datasets (genes expression matrix and 50 hallmark pathways score) was used to confirmed molecular subtypes. For confirming the appropriate numbers of molecular subtypes, we assessed the clustering prediction index (25) and Gaps statistics(26). We used the MOVICS package (including iClusterBayes, moCluster, CIMLR, IntNMF, ConsensusClustering, COCA, NEMO, PINSPlus, SNF, and LRA) with the two datasets to identify two specially molecular subtypes with different prognosis and silhouette score are similarity.

### Differentially expressed genes associated with molecular subtypes

We used the limma packages to confirmed differentially expressed genes of the two subtypes, and the P<0.01 and absolute fold-change > 1 was considered as statistical significance.

### Constructed the score

The pearson was used to confirm the correlation between the clusters and genes expression/50 hallmark pathways activity. And when the genes were positive with the clusters (cor>0.6) was identified as signature A and as negative with clusters (cor<-0.6) was confirmed as signature B. And then the ssgsea algorithm was used with 78 signature B genes to built a score which can effect quantify the two subtypes.

### External validation

The TIDE website (28) was used to predict immunotherapy effect based on the 535 LUAD patients. And also the Transcriptome data of IMvigor210 dataset (29) was downloaded which the patients was reviewed PD-1 blockade treatment. The two datasets above was used to conduct validated analysis.

### Statistical analysis

The R software (v4.10) was used to conducted all analyses. And we used the Wilcoxon test to analysis the significance between the two subtypes. The pearson correlation coefficient was used to assess the relationship between the two factors. All the results can be identified as statistically significant when P < 0.05.

## Result

### Confirmed two molecular subtypes

We used the MOVICS package (including iClusterBayes, moCluster, CIMLR, IntNMF, ConsensusClustering, COCA, NEMO, PINSPlus, SNF, and LRA) to conducted clustering with the two datasets (genes expression matrix and 50 hallmark pathways). Based on the results of CPI and Gaps analyses, we identified two molecular subtypes with different prognosis outcomes (figure1E) and the same score (figure1D). The distribution of multi-omics data in the two subtypes is shown in Figure1B.

**Figure1:**
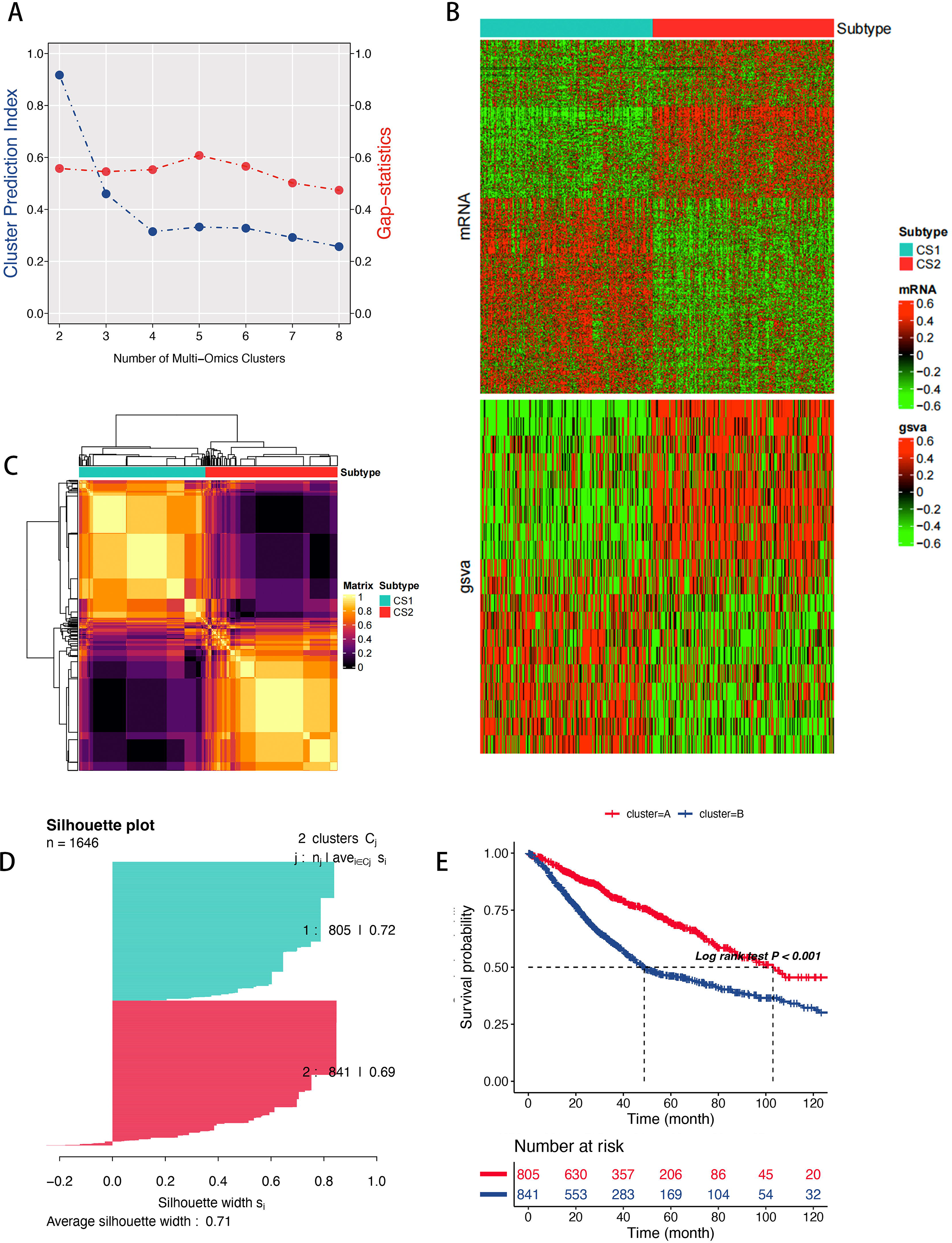
(A)The identified of clustering number by the CPI analysis and gap-statistical. (B) Consensus matrix based on the various algorithms. (C) The landscape of Various data of 11706 genes expression, 50 hallmark pathways in different molecular subtypes. (D) Silhouette-analysis evaluation. (E) survival analysis of the two clusters.

### Signatures of Molecular subtypes

We used the pearson to confirm the correlation between the clusters and genes expression/the tumorigenesis-related pathways score, and the cor>0.6 was identified as signature A and as the cor<-0.6 was confirmed as signature B (figure2A,B). And based on the signature A genes, we used the ssgsea algorithm to quantify the clusters. As the figure2C shown, the high score group had high rate cluster B and had high correlation with the tumorigenesis-related pathways (figureS1). The high score group showed better survival rate than the low score group(figure2F). Our results showed that the score model can better evaluate the worst prognosis outcome which is due to the high activity of tumorigenesis-related pathways such as E2F targets, G2M checkpoint, mtorc1 signaling, MYC targets V1 and MYC targets V2(figure2D,E). A total of 235 patients’mRNA expressions of GSE50081 and GSE3745 were used to be the external validation cohort and the results showed that the high score group had worst prognosis outcome and high activity of cell proliferation pathways (figure2G,H).

**Figure2:**
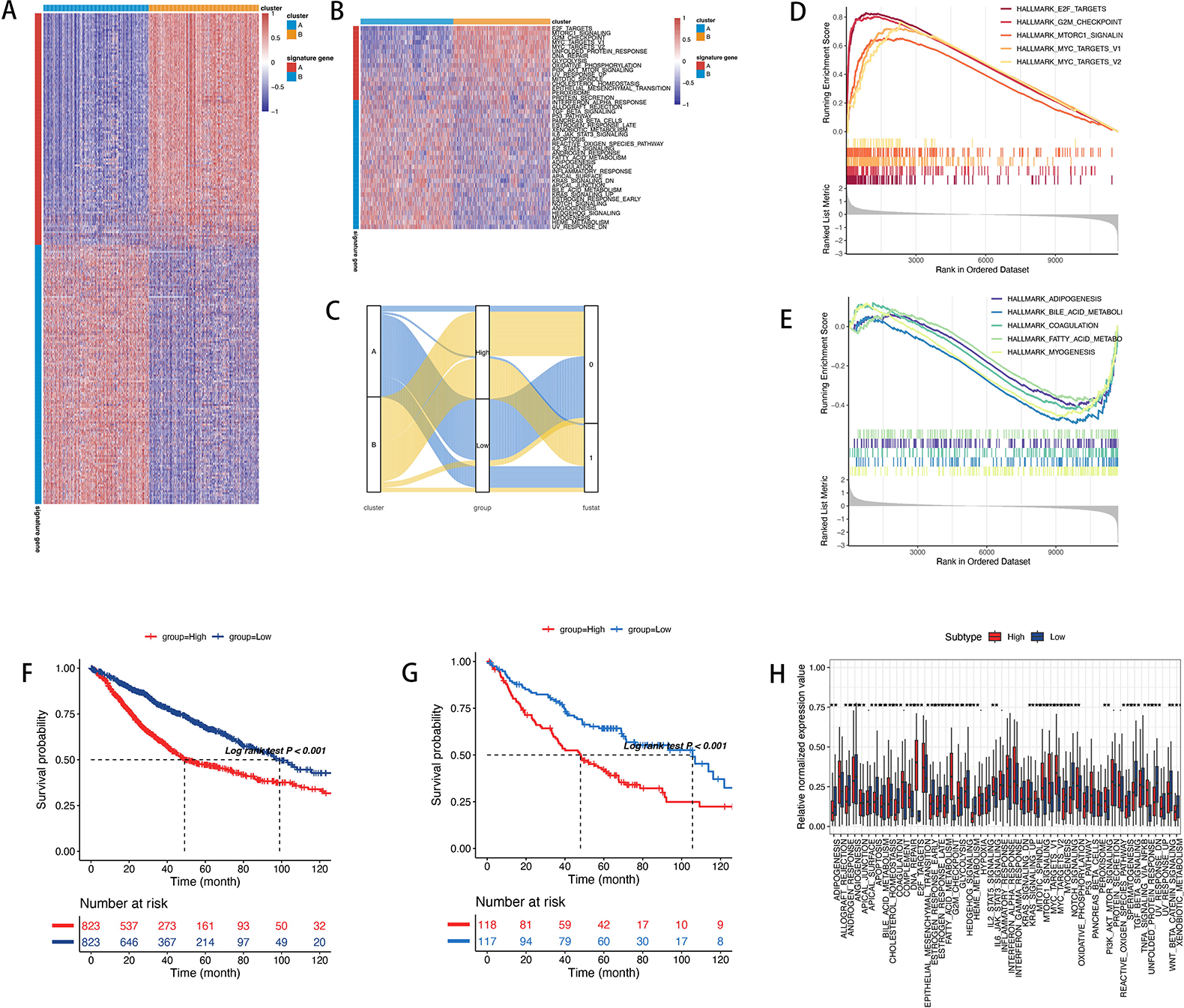
(A) Clustering analysis in DEGs of two clusters. (B)Clustering analysis in 50 tumorigenesis-related pathways of two clusters. (C)The distribution of clusters, and survival outcome in different score groups. (D,E) The biological function of low (E) and high score (D) groups. (F) Survival analysis of the two score groups. (G) Survival analysis of the two score groups in external cohorts (GSE50081 and GSE3745). (H) The landscape of 50 tumorigenesis-related pathways in the different score groups in external cohorts (GSE50081 and GSE3745).

### Genetic and epigenetic event between different score groups

A newly study shown that the genomic variation can be a source way which can drives tumor evolution and may provide some potential prognosis information. And a few of studies had addressed the prognostic value of CNA patterns in cancers. As the news study shown the CNA of 81 patients of esophageal squamous cell carcinoma and based on that 4 genes were find that had high correlation with the prognosis (10). Another study shown that gene CNA characteristic pattern had high correlation with prognosis in the breast cancer(11). In addition, the results from the classifiers based on the differences of CNA between LUAD and LUSC can help us deep the understanding of the carcinogenesis of lung cancer (12,13). There are differences of CNA lineage between LUAD and LUSC, and based on that we can explore the prognostic predictive value of CNA in LUAD subtypes.

As the figure3C shown that the high score group had high significant differential in the genomic variation. Also the high score group had instability genetic alteration, especially the copy number amplification of 8q24.2 and 14q13.3 (figure3D,E,F) (figureS2). For the amplification, we observed that the NKX2-8 and PAX9 located in the 14q13.3 was significant gain in the high score group than the low score group (S3). The high gain copy number of NKX2-8 and PAX9 may be the an important reason for the poor prognosis of high score group.

**Figure3:**
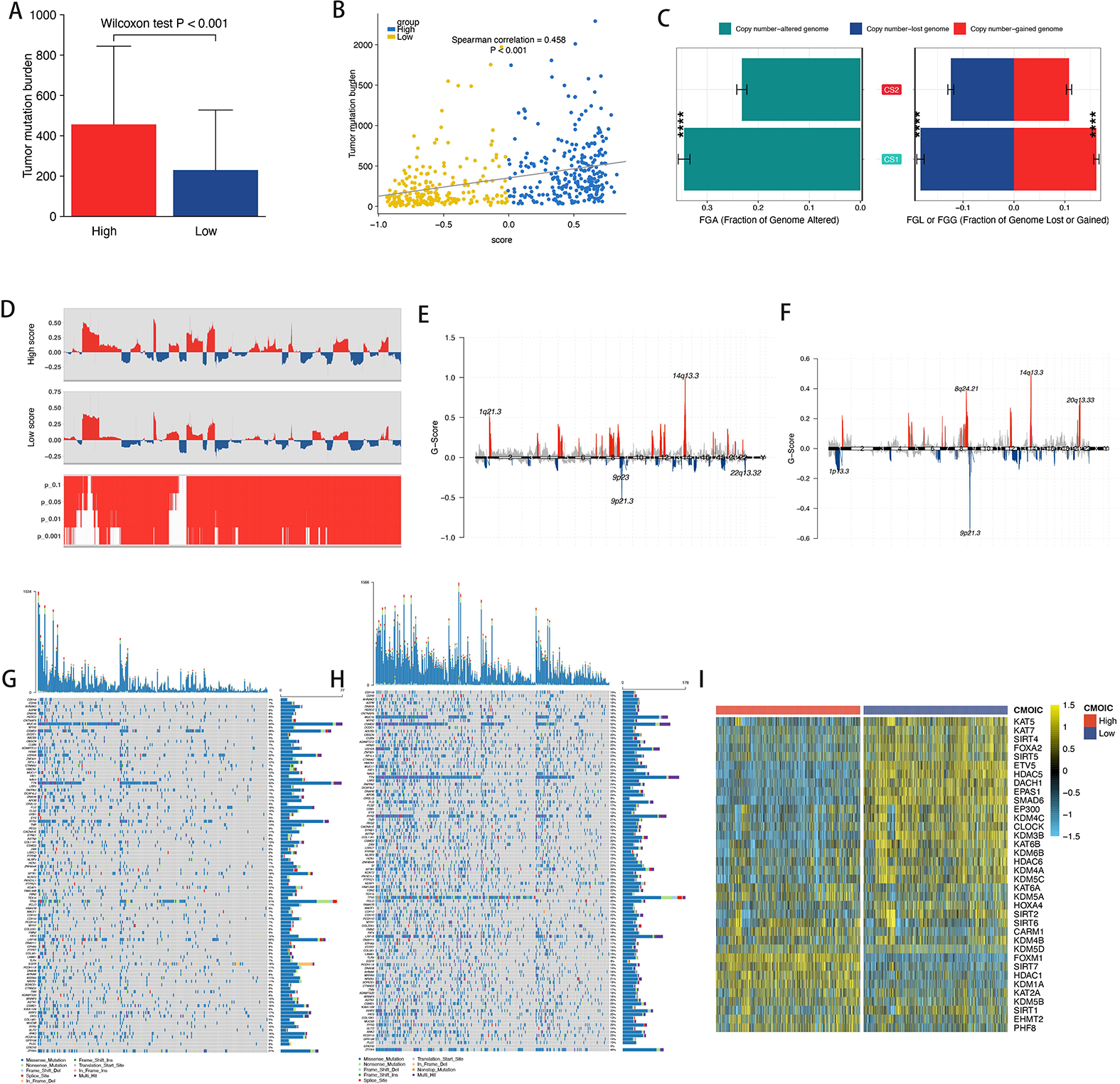
(A) The comparison of tumor burden mutational in different score groups. (B) The correlation between score and TMB. (C) The distribution of fraction genome altered (FGA) and fraction genome gain/loss (FGA/FGG) in the to different score group. (D) CNA plot showed the relative frequency of copy number gains (red) or deletions (blue) between the high score group and low score group of the LUAD cohort. (E,F)The distribution of the copy number of genomic regions between the high score group (E) and the low score group(F). (G,H) The significant mutation genes in the high scores (H) and low scores(G). (I) Heatmap showing profiles of regulon activity for 43 transcription factors including 8 LUAD-specific transcription factors and 35 chromatin remodeling potential regulons.

And we assessed the somatic mutations between different score groups of TCGA-LUAD cohort for presenting the genetic alterations. And the high score group showed that the high score group had higher TMB and more oncogene mutation than the low group(Wilcoxon test p < 0.001, Figure 3A, G, H). And the mutation frequency of oncogenes had significant differential between the different score groups, our study showed that the TP53/TTN had higher mutation frequency in the high score group than the low score group. The score had showed a high positively correlation with the tumor mutational burden(R = 0.458, p < 0.01)(figure3B). we used the MAFtools to analysis the distribution of significant genes mutation between the high score group and the low score group(figure3G,H). Above the results can help us to study the mechanism of the model and find a new target for targeted therapy. Also we assessed different activations of 8 LUAD-specific transcription factors and 35 potential regulators of chromatin remodeling. As shown in Figure 2I, the high score group was regulated by F0XM1, SIRT7, HDAC1, KDM1A, KAT2A, KDM5B, SIRT1, EHMT2 and PHF8. And the low score group had higher activity of KAT5, KAT7,SIRT4,FOXA2, SIRT5, ETV5, HDAC5, DACH1, EPAS1, SMAD6, EP300, KDM4C and CLOCK. Different information about the activity of regulators in the two score groups (recognition of epigenetically driven transcriptional networks) is an important distinction between the score groups.

### Immunotherapeutic Response

Our results showed that the high score presented high TMB, which can implied that the high score group had a huge potential to conducted immunotherapy especially the use of PD-1 or PD-L1 specific monoclonal antibodies. In our research, we used the datasets of TCGA cohort and IMvigor210 cohort to analyze the value of the score in the term of immunotherapy. We constructed the score with the signature A genes in the two datasets including the IMvigor210 cohort and TCGA cohort, and then assessed the effect of anti-PD-L1 immunotherapy in the two datasets. The high score group exhibited better objective effective rate of anti-PD-L1 treatment in the TCGA cohort and IMvigor210 cohort (figure4),which the high score had 12.9% response rate in the TCGA cohort and in the IMvigor210 cohort it had 30.2% response rate.

**Figure4.**
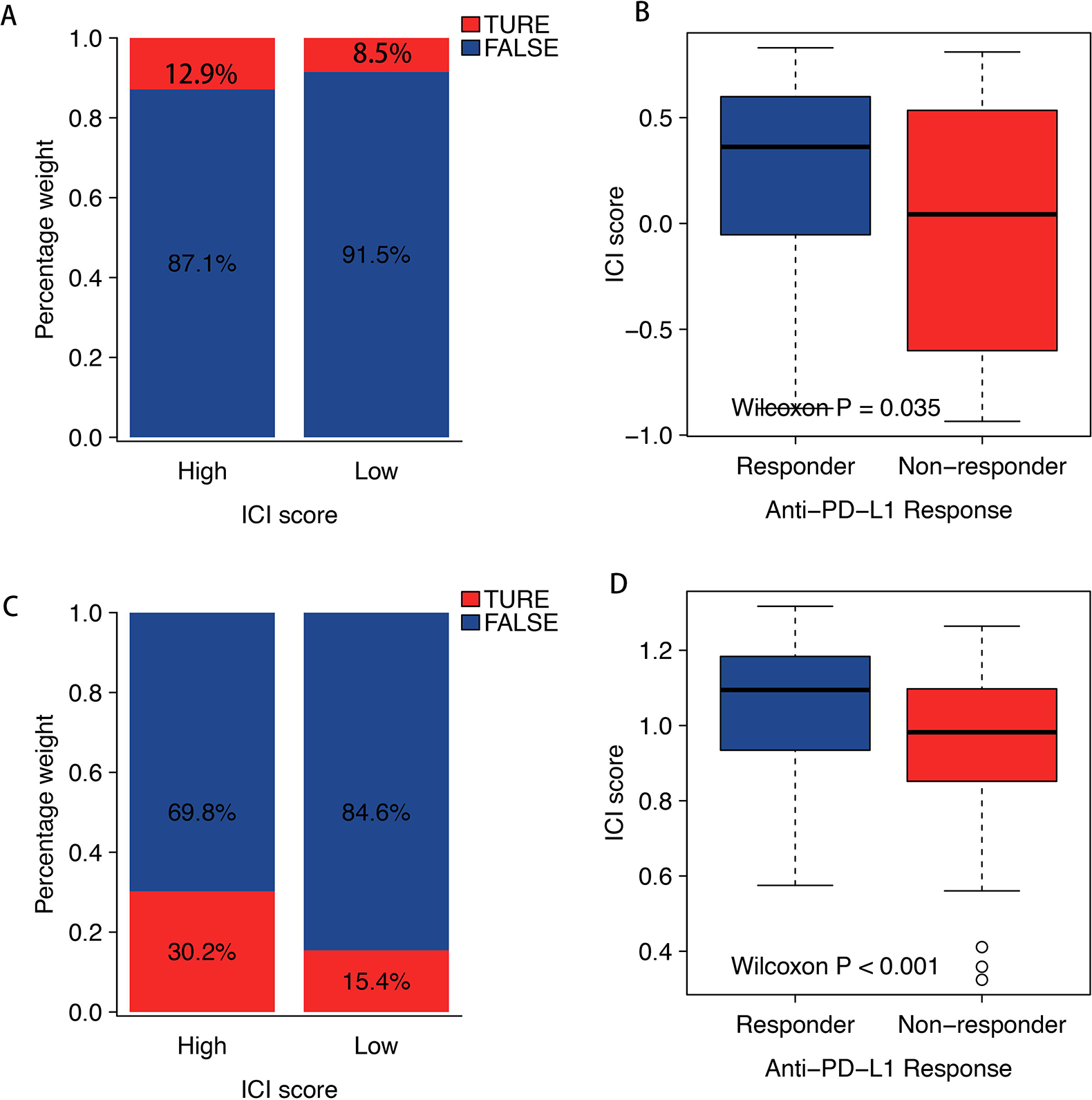
(A,C)The distribution of anti-PD-1 response rate in different score groups in the TCGA cohort and IMvigor210 cohort. (B,D) The two score groups with different anti-PD-1 response in the TCGA cohort and IMvigor210 cohort.

### The effective compounds of the two score groups

In order to confirmed the effective compounds of two score groups, the GDSC data was used to select. We selected the 78 signature A genes as the key genes of cluster B, and used the ssgsea algorithm to built the score withe lung cancer cells lines. Our results showed that the signature A genes had high significant expression in the mRNA levels in cells lines of lung cancer(figure5C). The high score group showed high significant sensitivity in PIM kinases (AZD1208), and the low score group may benefit from the p53 target therapy (NSC319726) (figure5D) (tableS4).

**Figure5.**
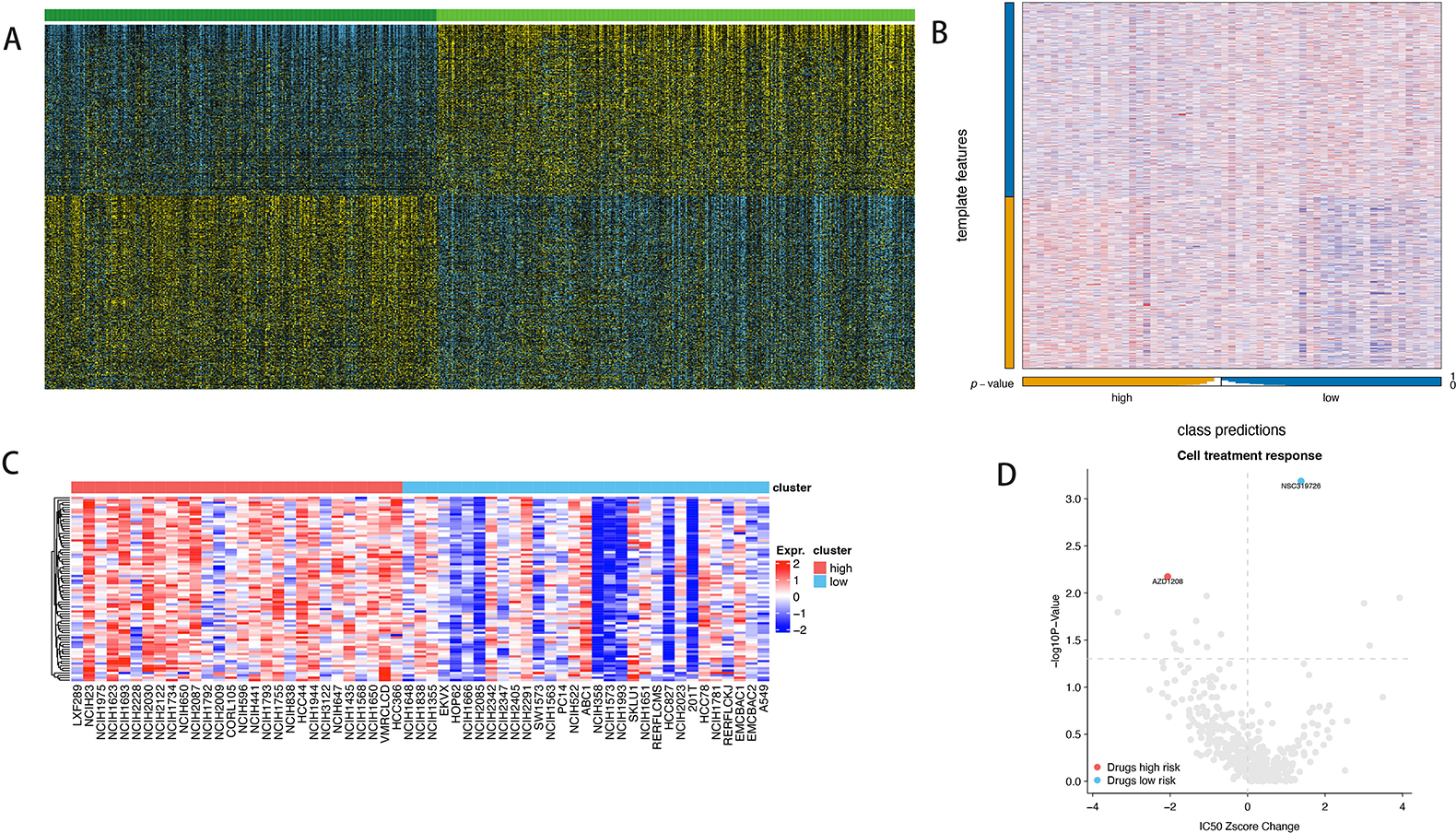
(A) Heatmap of subtype-specific upregulated biomarkers using limma for identifying the two score groups. (B) Heatmap of NTP. (C) Heatmap shows the expression pattern of 78 genes in cancer cell lines showing low or high score group. (D)The relative changes of the ic50z score and P values in selected luad cell systems at low or high score group.

### The ECT2 was identified as the Key Nodes in a Gene Regulatory Network for malignant phenotype

A total of 78 signature A genes are uploaded to the STRING database to build the PPI network (score_threshold=400). The PPI network is shown in the figure6A. The analysis of protein–protein interaction according to the STRING database and COX analysis highlighted ECT2 genes as key nodes with in the network. According to univariate cox regression analysis, the ECT2 was significantly correlated with overall survival (OS) (HR = 1.367, p-value < 0.01) (tableS1).

**Figure6.**
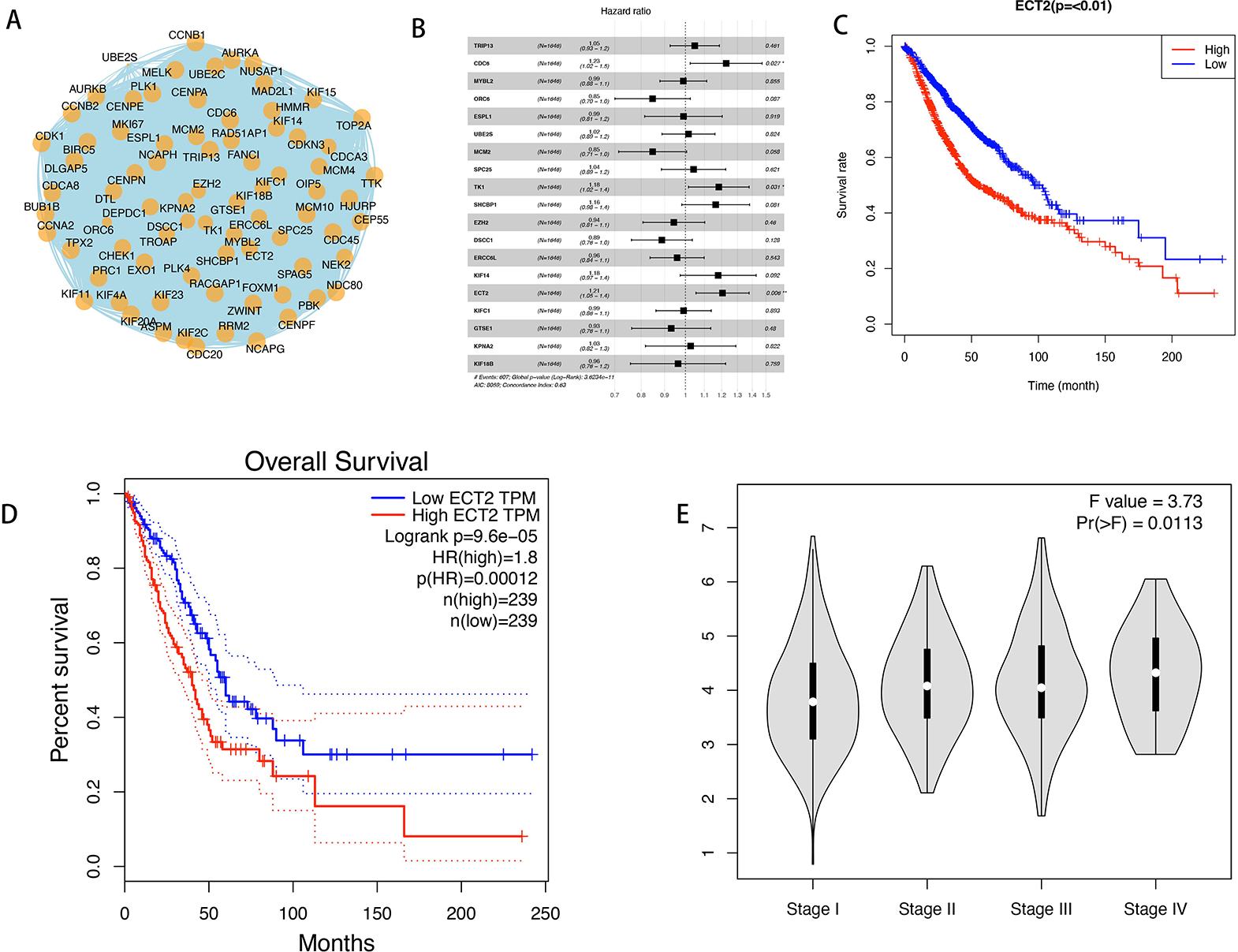
(A) PPI network of the signature A genes. (B)multivariate Cox regression analysis. (C)mRNA expressions of ECT2 in lung cancer and normal tissues as accessed by GEPIA database. (D)The overall survival analysis of ECT2 in the 1646 patient of lung cancer. (E)The overall survival analysis of ECT2 in the GEPIA analysis. (F)Box plot of the ECT2 for different pathological stages in the GEPIA analysis.

Meanwhile, according to multivariate Cox regression analysis, the ECT2 was an independent prognostic indicator (HR = 1.205, p-value < 0.01) (tableS2,Figure 6B) and our result showed that the ECT2 expression had high expression with the advance stage(figure6E).

## Discussion

Lung cancer is the mainly reason of cancer-related deaths worldwide, and was diagnosed about 2.21 million case and 1.80 million deaths in 2020(30). Now, the lung cancer patients had only 19% survival rate of 5 year survival rate, among which 57% was diagnosed as advanced stage(31). There are high rate of mortality and incidence, and so it important to detect in early times. Currently, the LDCT (low-dose computed tomography) was used widely to apply to screen the person who at the high risk of cancer. However, the false positives of LDCT would cause unnecessary and false treatment, and also add the cost and negative social and psychological consequences (32-34). Thus, it is urge to find an effective biomarker to help improve the screening methods and clinical treatment methods, and ultimately improve lung cancer prognosis. The multi-data analysis helps to classify molecular subtypes of LUAD which can precisely guide treatment choices and predict the prognosis. And the various gene expression was used to help us strength the understanding of LUAD biology and multiple molecular classifications. it is significant in the development of malignant phenotype that various different changes in molecular subtypes. The single-level histological is used to reveal the mechanisms of cancer development with high-throughput techniques in epigenomes, genome libraries, proteins, transcripts, microbiomes (35). And it is still no fully explain the complexity of the mechanisms of oncogenic driving and malignant phenotype in the single molecular-level approach. Therefore, a LUAD classification scheme based on multiomic characteristics is proposed, which may reveal the heterogeneity of LUAD.

Here, we used the GSVA algorithm to built 50 tumorigenesis-related pathways score. Based on the 50 tumorigenesis-related pathways score and genes expression matrix, the MOVICS algorithm was used to identified molecular subtypes. In our results, the cluster B had high activity of tumorigenesis-related pathways and poor prognosis outcome. And based on those DEGs/50 tumorigenesis-related pathways score, We used the pearson correlation to assess the relationship between genes/tumorigenesis-related pathways and clusters. And 78 genes and 4 tumorigenesis-related pathways score were identified as the cluster B mark genes/pathways(tableS3,S4). The ssgsea algorithm was used to construct score with the 78 signature A genes which can effect quantify the clusters, and the results showed that the high score group is high overlap with the cluster B. The high score group exhibited worst survival rate and the high activated of cell proliferation related-pathways, such as E2F targets, G2M checkpoint, mtorc1 signaling, MYC targets V1 and MYC targets V2. Also, we used two external cohort (GSE68465 and GSE72094) to verify our score accuracy in predicting the prognosis of lung cancer patients, and our results showed that the score exhibited a good predicting for prognosis. A somatic mutation analysis elucidated a higher frequency of TP53 and TTN mutations and low EGFR mutations in the high score group of TCGA-LUAD cohort. The mutation of TP53 can cause the loss of tumor suppressability and accelerates tumor formation (14). A newly study showed that the mutation of TP53 can influence the sensitivity of ICIs, regulate the expression of immune checkpoints, activate effector T cells, and also it reported that it affect a group of genes involved in cell cycle regulation, DNA replication, and damage repair in lung cancer [15-17]. The TTN had high mutation rate in various cancers and correlated with the the response to checkpoint blockade in solid tumors (18). Also, the detection of TTN mutations in peripheral blood is associated with satisfactory objective response and survival of ICBs (19). Recent studies have also shown that TTN mutations have great potential as predictive markers for ICBs in LUAD patients (20). In our study the high score had high TMB which may correlate with the higher frequency of TP53 and TTN mutations. Based on the high TMB of high score group, in our study, high group score can be identified as the most promising immunotherapy subtype. We used IMvigor210 cohort and TCGA cohort for corcorrelation with scores. The results showed that in both cohorts, the high group had a higher rate of anti-PD-L1 treatment than the low score group.

The PIM kinases worked as oncogenes which has a class of serine/threonine kinases, and can strength cell proliferation by transcriptional activation of genes (21). The PIM kinase has three isoforms including PIM-1, PIM-2, and PIM-3, and which plays an plays an important role in tumorigenesis, such as the PIM-1 is high expression in solid tumors and hematological tumor, the PIM-2 was reported that it is overexpressed in the lymphoma, myeloma and leukemia, also the PIM-3 had been confirmed it is highly expressed in adenocarcinoma. Newly study about PIM kinases showed it plays different roles in tumorigenesis, including multiple myeloma proliferation, anti-apoptosis, cell cycle regulation and mediating bone destruction (22,23). PIM kinase inhibits apoptosis in prostate cancer cells and promotes cell cycle progression, and its overexpression is related to fractionation and tumor transformation(24). In our study, the expression of PIM1 and PIM2 in the high score group are higher than the low score group(figureS4). We used the GDSC data to predict an effective compound for different score groups. And the results showed that as a PIM inhibitor--AZD1208 had high sensitivity in the high score group. And the a p53 (R175) mutant activator that inhibits the growth of cells -NSC319726 had high sensitivity in the low score group.

In a result, we used multi-omics datas to built a model to predict the prognosis of LUAD and the high score group can be identified as the high activity of cell proliferation and poor survival. And we used IMvigor210 cohort and TCGA cohort for corcorrelation with scores. The results showed that in both cohorts, the high group had a higher rate of anti-PD-L1 treatment than the low group which can be ascribed to higher frequency of TP53/TTN mutation and high tumor mutation burden.

In conclusion, our study built a score model to predict the activity of cell proliferation and prognosis, and a higher benefit from ICI therapy in LUAD with TP53/TTN mutation. Our study exhibits different cellular and molecular stratification of LUAD patients, and the high score group might benefit from the targeted inhibition of the PIM kinases and ICI therapy. Our study provide a series of evidence that that can indicate the specific alterations of epigenome prior in the different prognosis groups can improve the early stratification of patients with high invasion and proliferation. And also our study unraveled ECT2 expression as a characteristic feature of LUAD with a high activity of cell proliferation phenotype and as a promising drug target for combinatorial treatment strategies.

## Figure legend

FigureS1: The correlation between the score and the tumorigenesis-related pathways.

FigureS2: (A)Distribution of focal and broad copy number alterations among the different score groups. (B)Distribution of focal and broad copy number alterations in the 8q21.24 among the different score groups. (C)Distribution of focal and broad copy number alterations in the 14q13.3 among the different score groups.

FigureS3: (A)Distribution of focal and broad copy number alterations of NKX2-8 among the different score groups. (B)Distribution of focal and broad copy number alterations of PAX9 among the different score groups.

FigureS4: (A)The comparison of PIM1 expression between the different score groups. (B)The comparison of PIM2 expression between the different score groups

tableS1: Single variable Cox regression analysis

tableS2: Multivariate Cox regression analysis.

